# Temperate phage-antibiotic synergy is widespread, but varies by phage, host, and antibiotic pairing

**DOI:** 10.1101/2024.08.20.608816

**Authors:** Rabia Fatima, Alexander P. Hynes

**Author notes:** **Correspondence** Alexander P Hynes.

## Abstract

With a decline in antibiotic effectiveness, there is a renewed interest in bacteriophage (phage) therapy. Phages are bacterial-specific viruses that can be used alone or with antibiotics to reduce bacterial load. Most phages are unsuitable for therapy because they are ‘temperate’ and can integrate into the host genome, forming a lysogen which is protected from subsequent phage infections. However, integrated phages can be awakened by stressors such as antibiotics. This interaction was previously reported to result in a potent synergy between antibiotic classes and a model *E. coli* temperate phage, which can readily eradicate the bacterium at sub-lethal concentrations of antibiotics, despite the poor effectiveness of the phage alone. Here we explore the generalizability of this synergy to a clinically relevant pathogen: *Pseudomonas aeruginosa*. Thirty-six temperate phages isolated from clinical strains were screened for synergy with six antibiotics (ciprofloxacin, levofloxacin, meropenem, piperacillin, tobramycin, polymyxin B), using checkerboard assays. Interestingly, our screen identified phages that can synergize with each antibiotic, despite their widely differing targets - however, these are highly phage-antibiotic and phage-host pairing specific. Screening the strongest pairings across multiple clinical strains reveal that these phages can reduce the antibiotic minimum inhibitory concentration up to 32-fold, even in a resistant isolate, functionally re-sensitizing the bacterium to the antibiotic. When meropenem and tobramycin were effective synergistic agents, they did not reduce the frequency of lysogens, suggesting a mechanism of action independent of the temperate nature of the phages. In contrast, ciprofloxacin and piperacillin were able to reduce the frequency of lysogeny, the former by inducing phages – as previously reported in *E. coli*. Curiously, synergy with piperacillin reduced the frequency of lysogeny, but not by inducing the phages, and therefore likely acts by biasing the phage away from lysogeny in the initial infection. Overall, our findings indicate that temperate phages can act as adjuvants to antibiotics in clinically relevant pathogens, even in the presence of antibiotic resistance, thereby drastically expanding their therapeutic potential.

## Introduction

The widespread use of antibiotics has selected for resistance, resulting in a decline in their effectiveness and a rise in untreatable infections (1). As a result, there has been a renewed interest in treatments with alternative modes of action such as bacteriophage (phage) therapy. Phages are bacteria-specific viruses (2) which hijack the host cell machinery and redirect it to synthesize phage components, resulting in host cell lysis and release of new infectious phage progeny (3). These can also synergize with multiple classes of antibiotics, known as “Phage-antibiotic synergy” (PAS), which is associated with changes in phage replication and increase in phage production (4–7).

Most phage therapy work to date has been with ‘virulent’ (strictly lytic) phages. In contrast with these, **temperate phages** can undergo an additional replication cycle known as lysogeny a dormant state involving the integration of the phage genome into the bacterial host and replication along with it (8). The integrated phage is referred to as a **prophage** and a bacterium carrying a prophage is a **lysogen**. In addition to choosing between lysis and lysogeny at the time of infection, prophages are capable of exiting lysogeny and switching to lytic replication in response to external stressors in a process known as **induction** (8). This decision between lysis and integration, at the initial time of infection or later during dormancy, is facilitated by phage encoded proteins in many well studied temperate phage models (e.g. cI repressor in *Escherichia coli* phage Lambda) (9, 10). Even though this switch is genetically encoded, much of this decision is responsive to environmental factors (11–15). Of these, one of the most well known are antibiotics that trigger the bacterial SOS response (e.g. mitomycin C, fluoroquinolones, some beta-lactams) and result in phage induction through subsequent cleavage of the phage repressor protein (16–22).

Temperate phages have been overlooked for use in phage therapy because they present concerns of overgrowth of phage resistant lysogens because of immunity; superinfection immunity (23, 24), mediated by the phage repressor protein, or superinfection exclusion, mediated by phage proteins that block DNA-entry (25). However, due to the narrow host range of many phages, in instances when lytic phages are difficult to find (26, 27), studies have had to employ virulent variants of temperate phages obtained through genetic engineering for treatment of bacterial infections (28, 29). Interestingly, up to 75% of bacteria already contain a prophage in their genome (30, 31), greatly facilitating their discovery. The isolation and use of virulent mutants of temperate phages that can infect a lysogenic host has also been proposed for *P. aeruginosa* infections (32). These can also be of particular interest for pathogens such as *Clostridioides difficle* where strictly lytic phages have not been identified to date (26).

Studies examining the potential of temperate phages in therapy have been few and far between. Temperate phages of *Burkholderia cepacia* complex show synergistic interaction with other phages where their therapeutic potential inversely correlates with the frequency of lysogeny (33). In addition, temperate phages isolated from clinical strains of *Pseudomonas aeruginosa* were shown decrease twitching motility, important for virulence, in lysogens (34). Phage administration was shown to reduce bacterial load in mice and *Drosophila* model of *Pseudomonas* infection (35). Administration of a temperate phage was able to reduce bacterial load and prevent toxin production in an *in vitro* human colon model of *C. difficle* infection (36). However, the study also reported increased spore formation with potential for increased risk of re-emerging infection. Several studies have also proposed the use of temperate phage cocktails for *C. difficle* (37, 38). In a hamster model, Nale et al. (2016) reported that phage cocktail not only reduced the colonization load of *C. difficle* but could also delay symptoms by 33 h (38).

While temperate phages are currently not ideal for monotherapy, the use of adjuvants that can bias their decision away from lysogeny and towards lytic replication can be promising for compassionate last resort cases. A four temperate phage cocktail combined with Ca2+ or Zn2+ reduced methicillin-resistant *S. aureus* load by 2.64-fold compared to phage cocktail alone in a mouse model, however their frequency of lysogeny was not reported (39). Knezevic et al. (2013) demonstrated that temperate phage σ-1 combined with ¼ minimum inhibitory concentration (MIC) ceftriaxone reduced *P. aeruginosa* counts by ≥ 2 logs (41). The same effect was not observed with ciprofloxacin, a fluoroquinolone, gentamicin, a protein synthesis inhibitor, and polymyxin B, an outer membrane targeting antibiotic. The study briefly noted a potential involvement of antibiotic mediated phage induction.

Supported by the idea that SOS-response inducing antibiotics can act as phage inducers, we previously demonstrated that combination of temperate phage HK97 and sublethal ciprofloxacin, can synergistically reduce *E. coli* survivor count up to 10^8^-fold after an 18 h treatment, largely by inducing any lysogens that formed. This was dubbed temperate phage antibiotic synergy (tPAS) (42). This was generalizable to other antibiotics, including quinolones, anti-folates, and mitomycin C – all known to induce the bacterial SOS response (43). Interestingly, protein-synthesis inhibitors of several classes also show comparable synergy in this model, although these were determined to act by biasing the phage away from lysogeny during the initial infection, rather than by inducing lysogens (43). This is further supported by the finding that the potential of temperate phages in the clinical

With the reported efficacy of tPAS in *E. coli* demonstrating substantial reduction in lysogeny, the major concern in the therapeutic use of temperate phages, here we systematically investigate its effectiveness across phages, hosts, and antibiotics in the clinically relevant pathogen *P. aeruginosa*.

## Results/Discussion

### Temperate phages which synergize with ciprofloxacin are readily isolated from clinical strains

To establish tPAS in *P. aeruginosa*, we isolated 36 temperate phages that could infect the strain *P. aeruginosa* PA14 from filtrates of overnight cultures of 191 *P. aeruginosa* clinical isolates, highlighting the abundance and ease of isolation of these phages (Fig. 1a). This is consistent with many reports of isolation of temperate phages from clinical strains of *P. aeruginosa and C. difficle*, often aided by mitomycin C (44–46).

**Figure 1.**
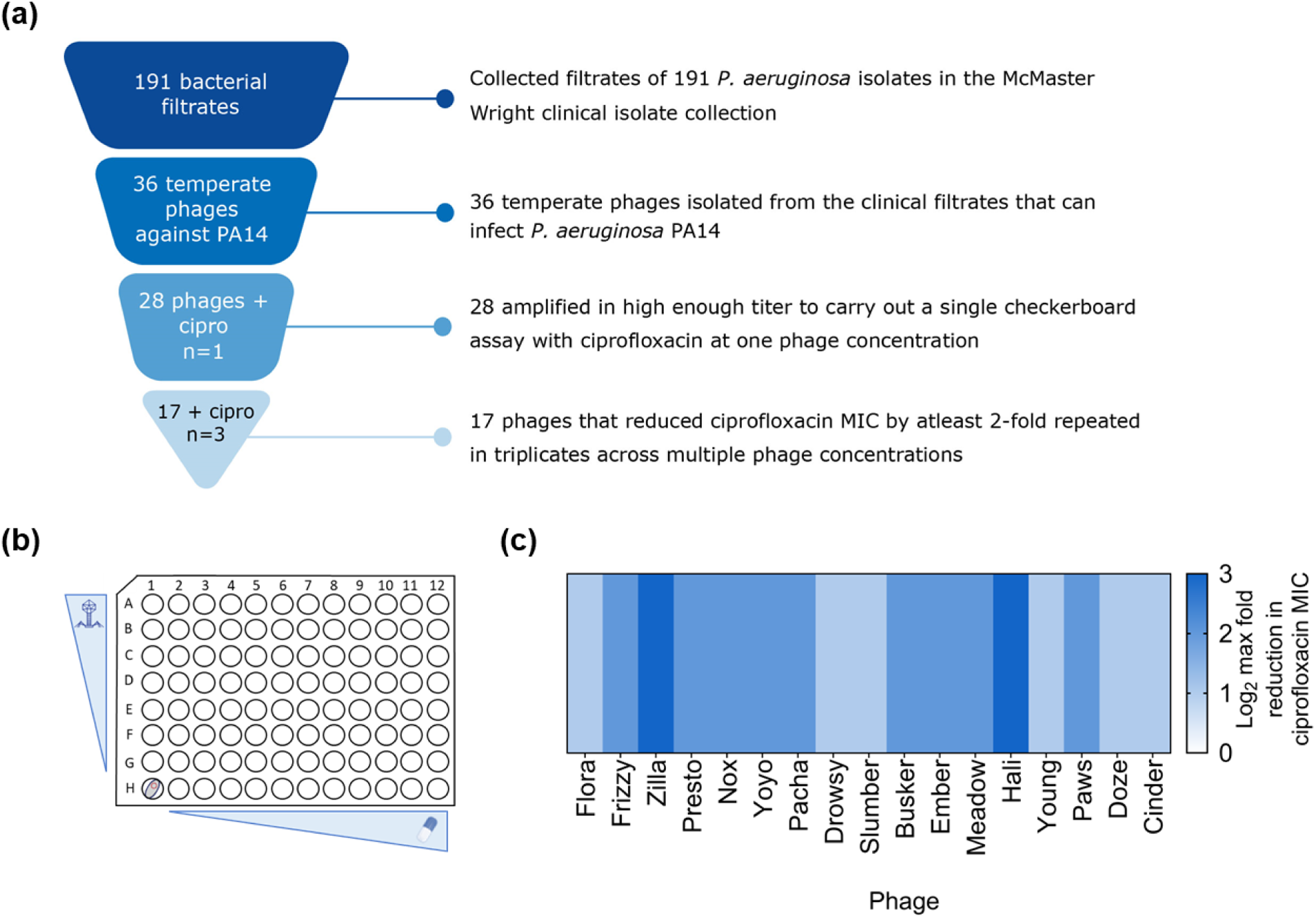
Temperate phages that synergize with ciprofloxacin can readily be isolated from clinical strains. **(a)** Workflow of isolation of temperate phages infecting *P. aeruginosa* PA14 from clinical strain collection and screening for synergy with ciprofloxacin in PA14. **(b)** Illustrative representation of checkerboard assay screening for synergy between a temperate phage and antibiotic. **(c)** Log_2_ maximum reduction in ciprofloxacin MIC achieved with the addition of a panel of temperate phages relative to no phage added control against *P. aeruginosa* PA14, plotted as a heat map (n=3 biological replicate).

Twenty-eight phages amplified on *P. aeruginosa* PA14 in high enough titre to screen for synergy with ciprofloxacin using a checkerboard assay in a single replicate (Fig. 1a-b). Seventeen phages that reduced the antibiotic MIC by at least 2-fold (limit of detection) were then repeated in biological triplicates. A summary of the screening with ciprofloxacin is shown in Fig. 1c as maximum fold reduction in the ciprofloxacin MIC achieved with the addition of the phage, regardless of the phage dose. All seventeen phages reduced the ciprofloxacin MIC by at least 2-fold, with the highest (8-fold) reduction achieved with phage Zilla and phage Hali. These findings highlight that synergy between temperate phage and antibiotics can be achieved in *P. aeruginosa* with multiple phages, at least with ciprofloxacin, a known bacterial DNA-damaging phage inducer.

The seventeen phages that exhibited a synergistic interaction with ciprofloxacin were further characterized using whole genome sequencing and bacterial host range analysis. Recognizable phage genomes were obtained for all phages except phage Drowsy. Genome annotation predicted a transposase in 15 out of the 16 phages (not present in phage Flora) and these phages cluster closely with other *Pseudomonas* transposable temperate phages (Fig. 2a). Host range analysis carried out using 98 *P. aeruginosa* clinical isolates reveals that these phages exhibit broad host range, with phage Busker able to infect 31 out of the 98 strains tested (Fig. 2b). While phage Drowsy and phage Slumber exhibited a similar host range, the efficiency of plaquing for four out twenty-nine strains differed. Despite variations in host ranges – some of which can presumably be attributed to host-controlled variation, based on sequence similarity and phylogenetic analysis of phages Yoyo, Pacha, Slumber, Busker and Ember, phage Ember was kept as a representative of that cluster.

**Figure 2.**
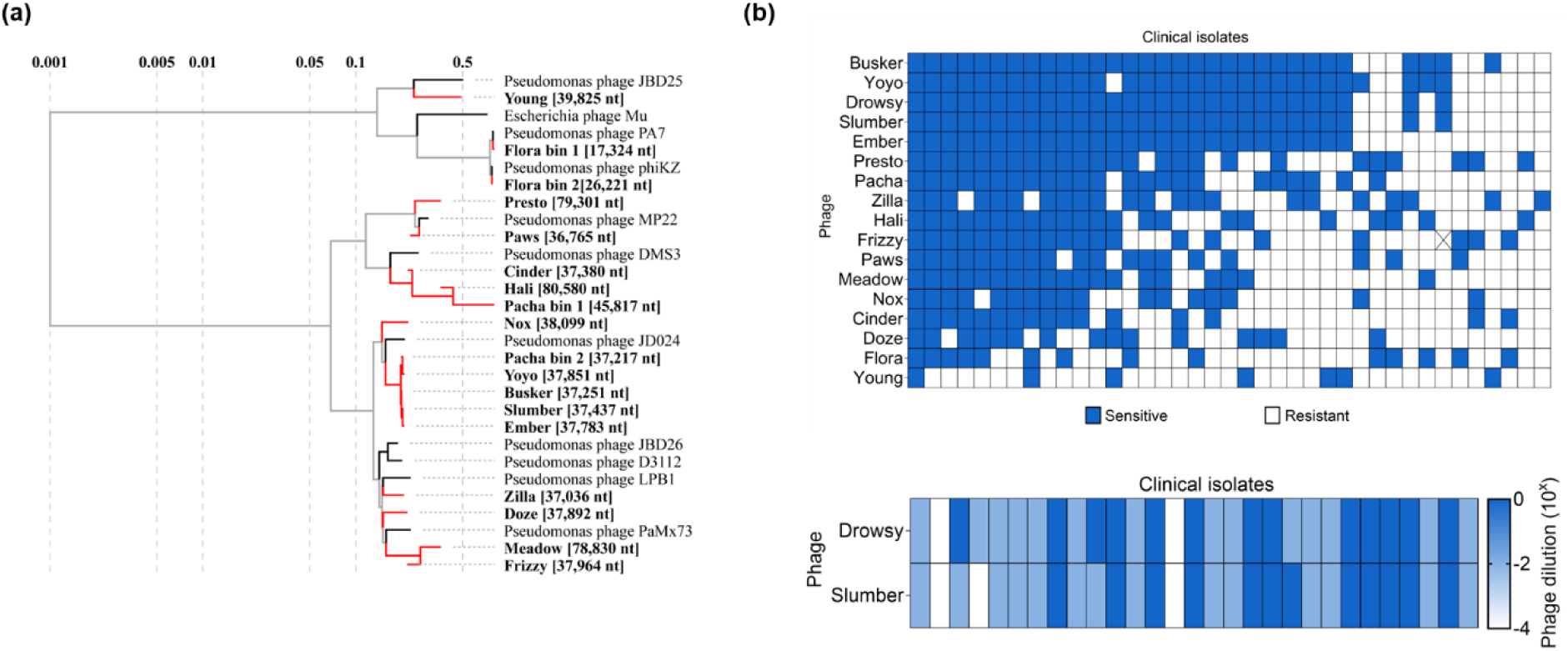
Newly isolated PA14 temperate phages are unique. **(a)** Phylogenetic tree of the isolated temperate phages. Tree was generated using ViPTree with a subset of related genomes. Branch lengths are indicated on a log scale. **(c)** Host range of the newly isolated PA14 temperate phages tested against 96 *P. aeruginosa* clinical strains. Each row represents a phage, ordered in largest to smallest host range, and columns represent a single isolate, ordered from susceptible to most to least phages. Blue denotes phage susceptible. **(d)** Host range comparison of phage Drowsy and phage Slumber. Heat map depicts the 10-fold phage dilution on which plaquing was observed on a specific host, represented by the columns.

### tPAS with ciprofloxacin results in bacterial eradication through induction

While checkerboard assays do establish a clear synergistic effect revealed as a reduction of the antibiotic MIC, they do not provide enough resolution to quantify this effect and determine the mechanism of bacterial growth inhibition. To better quantify this synergy over a range of antibiotic concentrations, we challenged *P. aeruginosa* PA14 with phage Hali and ciprofloxacin in liquid media. Survivors present after an 18 h challenge were plated with no selection for another 18 h. Fold reduction in survivor count relative to the untreated host was calculated for each challenge. There was a less than 5-fold reduction in survivor count when challenged with Hali alone and a more pronounced dose dependent ~10^4^ fold reduction with antibiotic at the highest concentration (Fig. 3a). These results highlight the poor efficacy of the phage and antibiotic alone in eradicating PA14.

**Figure 3.**
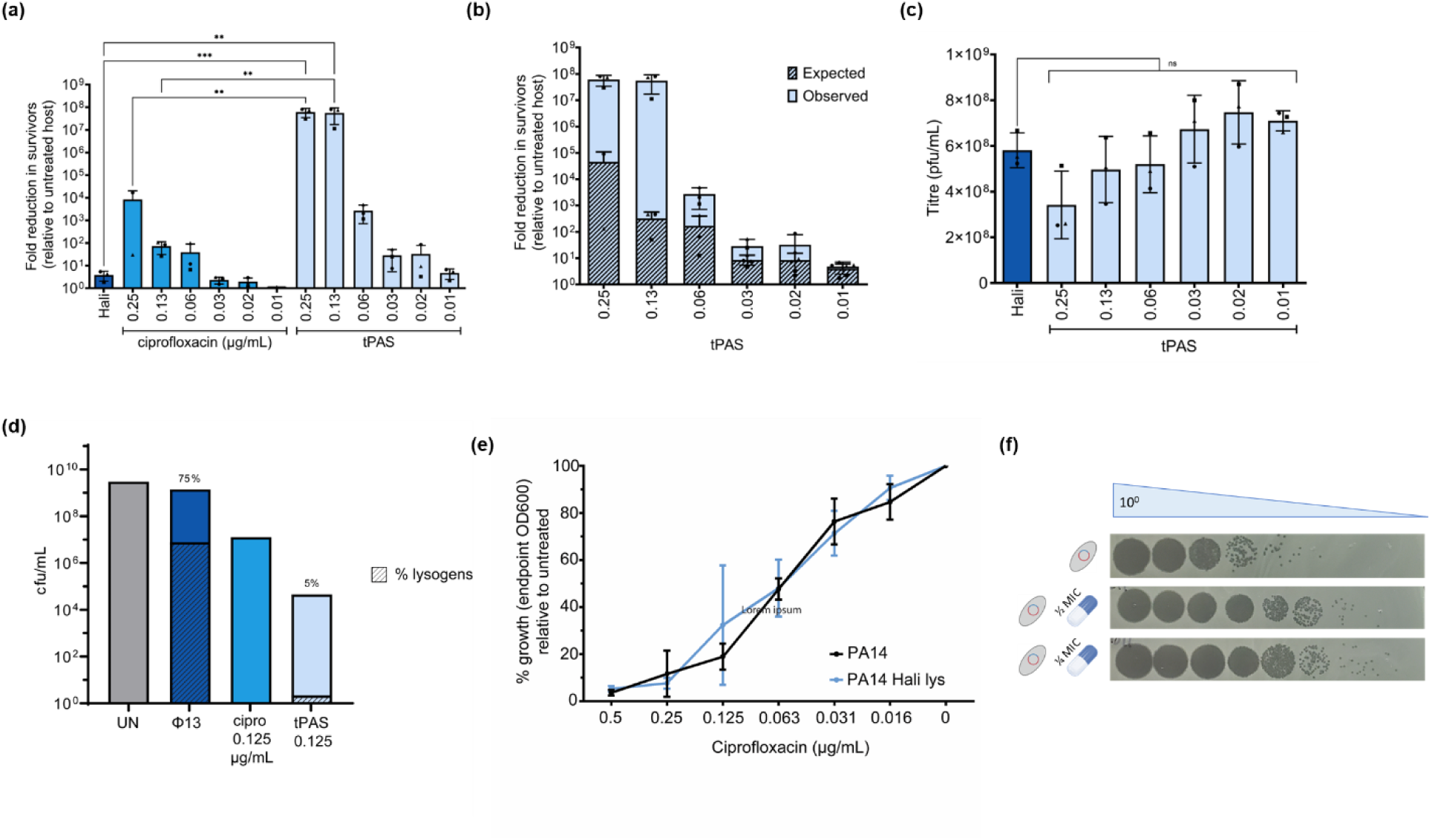
Phage Hali and ciprofloxacin challenge synergize to eradicate *Pseudomonas* through induction. **(a)** Fold reduction in PA14 survivors relative to untreated host (mean ± SD). In the case that no survivors were detected, the data were set to the limit of detection (1 colony). Each point indicates a biological replicate, denoted by the different shapes, counted in technical triplicates. **(b)** Observed fold reduction in survivors (solid) versus the expected (diagonal line) effect. Expected effect was calculated by multiplying the phage alone reduction with antibiotic reduction for the corresponding antibiotic concentration. **(c)** Phage quantification from the overnight challenges. Bars indicate average plaque forming units (PFU/mL) ± SD. Each point indicates a biological replicate, denoted by the different shapes, counted in technical triplicates. **(d)** Survivor quantification (cfu/mL) of untreated PA14, challenged with phage Hali ± ciprofloxacin at 0.125 ug/mL where the strongest synergy was observed in a repeat experiment. Striped bars indicate percentage of lysogens as determined using a lysogen stamp test, values also indicated above bar. Twenty survivors of phage ± ciprofloxacin were streaked purified for lysogen testing. The height of the % lysogen bar is proportional to the height of the total cfu/mL bar. **(e)** Ciprofloxacin MIC curve of PA14 and PA14 phage Hali lysogen represented as percent growth (endpoint OD600) relative to untreated host, plotted as mean ± SD (n=3 biological replicates, each in three technical triplicates). **(f)** Representative phage quantification of PA14 phage 13 lysogen challenged with ½ and 1/4 MIC ciprofloxacin.

In contrast, PA14 challenged with phage Hali in the presence of ciprofloxacin at the two highest concentration resulted in approximately a 10^8^-fold reduction in survivors, corresponding to a complete eradication. The number of survivors then decreased in a dose dependent manner as the antibiotic concentration decreased. However, the clearest evidence for a synergistic effect is obtained by comparing the observed data to the expected multiplication of the independent effects of phage and antibiotic (Fig. 3b). Phage Hali and ciprofloxacin resulted in the strongest synergistic effect at sublethal 0.13 µg/mL with an observed reduction that is roughly 10^5^ fold higher than the expected effect. This effect is much higher than that observed with temperate phage σ-1 and ceftriaxone in *P. aeruginosa* ATTC 9027 (40). However, it agrees with our previous work performed in *E. coli* with model temperate phage HK97 and ciprofloxacin (42), although across a smaller range of antibiotic concentrations.

To elucidate the mechanism through which phage Hali-ciprofloxacin pairing results in bacterial killing, we quantified phages in the filtrates of the phage-alone challenge and tPAS challenges from the overnight survivor quantification assay (Fig. 3c). We observed no significant difference in phage titre in the presence of the antibiotic compared to the phage alone challenge suggesting that phage Hali-ciprofloxacin synergy is not a result of a substantial increase in phage replication.

To determine if tPAS was instead biasing the phage lysis-lysogeny decision, we repeated the assay across a shorter range of antibiotic concentrations, purifying survivors from phage alone and a single antibiotic concentration where we observed the strongest synergy with regards to reduction in survivor count (Fig. 3d). Phage alone resulted in 75% lysogens, which reduced to 5% in presence of ciprofloxacin. The interaction works at the level of biasing the phage lysis-lysogeny equilibrium, potentially at the level of induction since antibiotics are well known to result in induction.

To confirm if PA14 lysogen of phage Hali can be induced with sublethal ciprofloxacin, we challenged wild type and lysogen with ciprofloxacin and quantified phages in the filtrates. Unlike in prior *E. coli* phage HK97 work (42), the PA14 phage Hali lysogen is not more sensitive to ciprofloxacin than the wild type (Fig. 3e), despite the roughly 100-fold increase in phage titer, relative to baseline spontaneous induction, in filtrates of lysogen grown with sublethal ciprofloxacin (Fig. 3f). Overall, our results indicate phage Hali and ciprofloxacin synergy results in bacterial killing through preventing the expansion of lysogen colonies likely through induction, where lysogens form but are subsequently induced in presence of sublethal antibiotic, like that reported in *E. coli* with phage HK97 and ciprofloxacin (42).

### tPAS can be achieved even with delayed administration of either agent

With a clear synergistic interaction observed with the co-administration of phage Hali and ciprofloxacin, we sought to investigate whether both agents need to be simultaneously present to achieve synergy. To first test if the antibiotic needs to be present when the phage infects, we monitored bacterial growth after simultaneous phage and antibiotic administration (Fig. 4a) or delayed antibiotic treatment by 4 h where the antibiotic likely only has lysogens to interact with (Fig. 4b).

**Figure 4.**
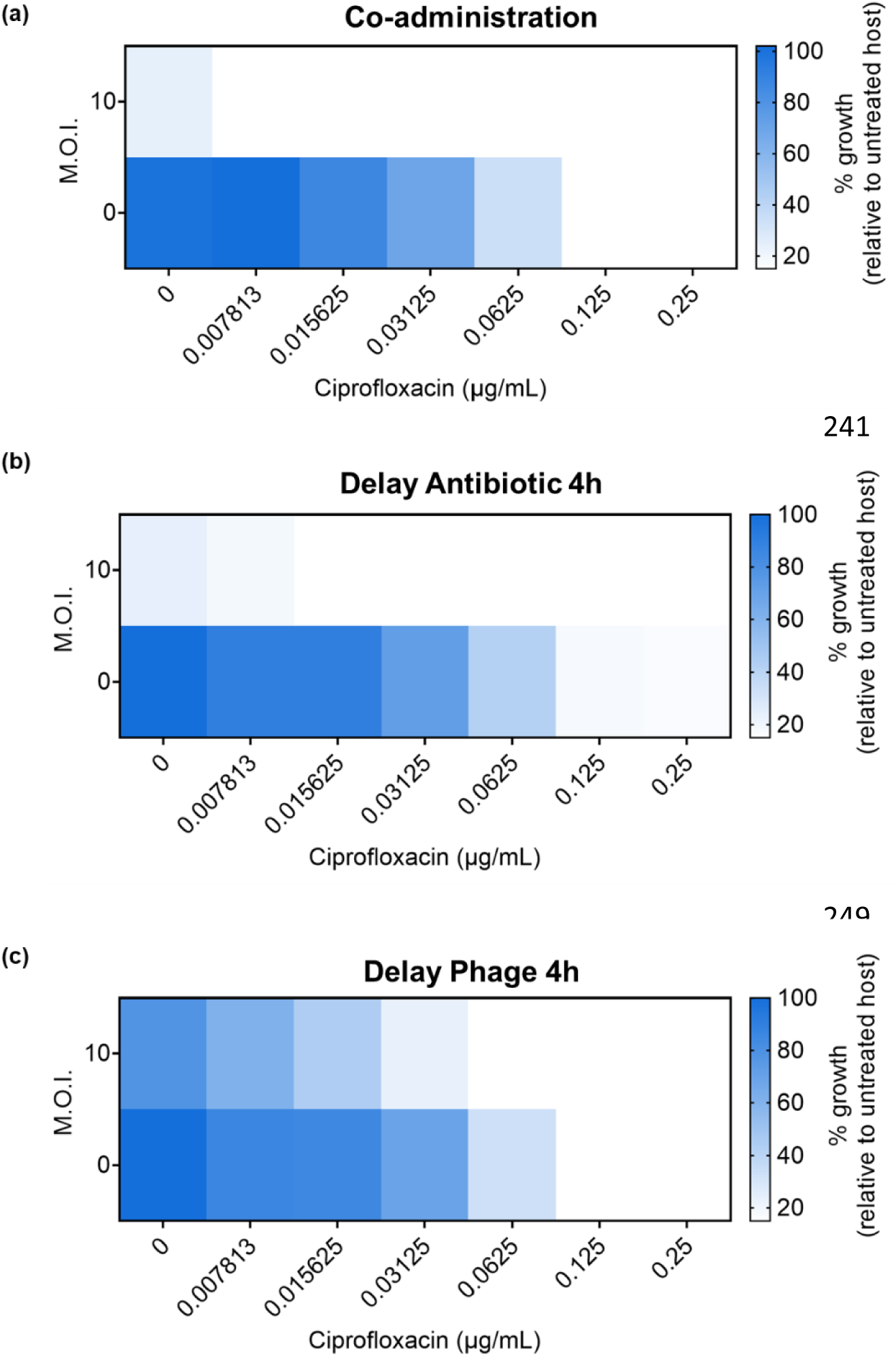
Temperate PAS can be achieved delaying antibiotic or phage. Checkerboard assay of phage Hali and ciprofloxacin in PA14 with **(a)** simultaneous, **(b)** delayed antibiotic by 4h, or **(c)** delayed phage by 4 h. Endpoint growth (OD600) relative to untreated host, plotted as a heatmap (n=3 biological replicates, each in three technical triplicates).

Delaying the antibiotic by 4 h results in an increase in antibiotic MIC compared to administering it alone at time zero (Fig. 4b). However, a comparable level of synergistic reduction in ciprofloxacin MIC was still observed in the presence of phage Hali when the antibiotic was delayed. Ciprofloxacin does not necessarily need to be present when the phage infects to achieve the same levels of tPAS as co-administration. Taken together with the frequency of lysogeny (Fig. 3d), the results highlight that phage Hali and ciprofloxacin synergy works through biasing lysis-lysogeny decision at the level of induction.

We also tested the effects of delaying the phage to investigate if pre-stressing the bacteria with antibiotic prior to phage treatment would result in synergy. Delaying the phage by 4 h results in only a 2-fold reduction in ciprofloxacin MIC (Fig. 4c). While the antibiotic does not necessarily need to be present at the time of phage infection, the opposite is not true. By delaying the phage, we drastically reduce the range of antibiotic concentration at which tPAS is observed.

It is important to note that the efficacy of delayed administration is likely dependent on the phage-antibiotic pairing. *P. aeruginosa* virulent phage JG024 demonstrates synergy with ciprofloxacin even with 1 h delay of either agent (47). The effect is lost if the treatment occurs after 6 h. In contrast, with ceftriaxone, only delaying phage JG024 by 1 h can inhibit bacterial growth and delayed antibiotic has no effect. Delaying ciprofloxacin (8 x MIC) and gentamicin (1 x MIC and 8 x MIC) 6 h post lytic phage treatment also reduced viable cell count >10^2^ cfu/mL (limit of detection), better than sequential treatment, in mono species *P. aeruginosa* 48 h biofilm (48). With regards to tPAS, the extent to which one of the agents can be delayed would presumably depend on the rate at which lysogens form in a specific host.

### tPAS is generalizable across phage-antibiotic pairings

Having established this interaction with ciprofloxacin, we sought to test the generalizability across antibiotics, covering three more antibiotic classes in addition to fluroquinolones (Fig. 6). These antibiotic classes were selected for their reported ability to either induce phages or their clinical relevance as anti-pseudomonal drugs. Beta-lactams are cell wall synthesis inhibitors that bind to penicillin-binding protein (PBP) to prevent crosslinking (49). While these have been shown to result in phage induction in multiple bacterial hosts (21, 50), the exact mechanism through which they induce phages, directly or indirectly through the bacterial SOS response (51–53), remains unclear. The beta-lactams, meropenem and piperacillin are commonly used in the clinic to treat *Pseudomonas* infection, where the latter is combined with a beta-lactamase inhibitor, tazobactam (54).

Additionally, aminoglycosides inhibit protein synthesis through binding to the A site of 16S ribosomal RNA, inhibiting translocation resulting in protein mistranslation (55). Since phages need host protein machinery for replication, protein synthesis inhibitors are not expected to synergize with antibiotics. Kanamycin antagonizes replication of *E. coli* phage T3, demonstrated as decrease in efficiency of plaquing, bacterial growth, and biofilm biomass, by decreasing phage burst size independent of changes in phage adsorption (56). Similarly, both kanamycin and apramycin reduced efficiency of plaquing of temperate phage Lambda by up to 1000-fold (57). Select protein synthesis inhibitors were also reported to antagonize *P. aeruginosa, S. aureus, and Enterococcus faecium* phages (58). Contrary to these studies, gentamicin and kanamycin both synergize with phage HK97 in *E. coli* through biasing the initial phage decision towards lysis at the time of infection (43). To investigate if a similar synergy between aminoglycoside and temperate phages can be observed in *P. aeruginosa,* we performed our assays with tobramycin, frequently used as an inhaled treatment for cystic fibrosis patients (59). Our screen also includes polymyxin B, an older outer membrane disrupting antibiotic which works through binding to lipopolysaccharide (60), that has regained interest as one of the last resort drugs for severe infections.

We observed synergy with all antibiotics, despite their vastly different bacterial targets, with most of the phages working particularly well in combination with piperacillin (Fig. 5). While some of these antibiotics have not been previously reported to interact specifically with temperate phages, strong synergy between virulent phages and several cell wall synthesis inhibitors (16/25 antibiotics tested), including piperacillin and meropenem, and protein synthesis inhibitors (5/25 antibiotics tested), including tobramycin, has been previously reported in *P. aeruginosa* (61, 62). Polymyxin B has also been reported to work in combination with virulent phages for treating *S. aureus* (63). However, we did not observe a consistent synergistic pattern with antibiotics of the same drug class, indicating that PAS here is phage-antibiotic pairing specific, previously also reported with a lytic phage and several antibiotic classes in extraintestinal pathogenic *E. coli* (7). Nonetheless, this drastically expands the library of compounds that can be used in combination with temperate phages for therapeutic use, even with antibiotics not previously known to induce phages.

**Figure 5.**
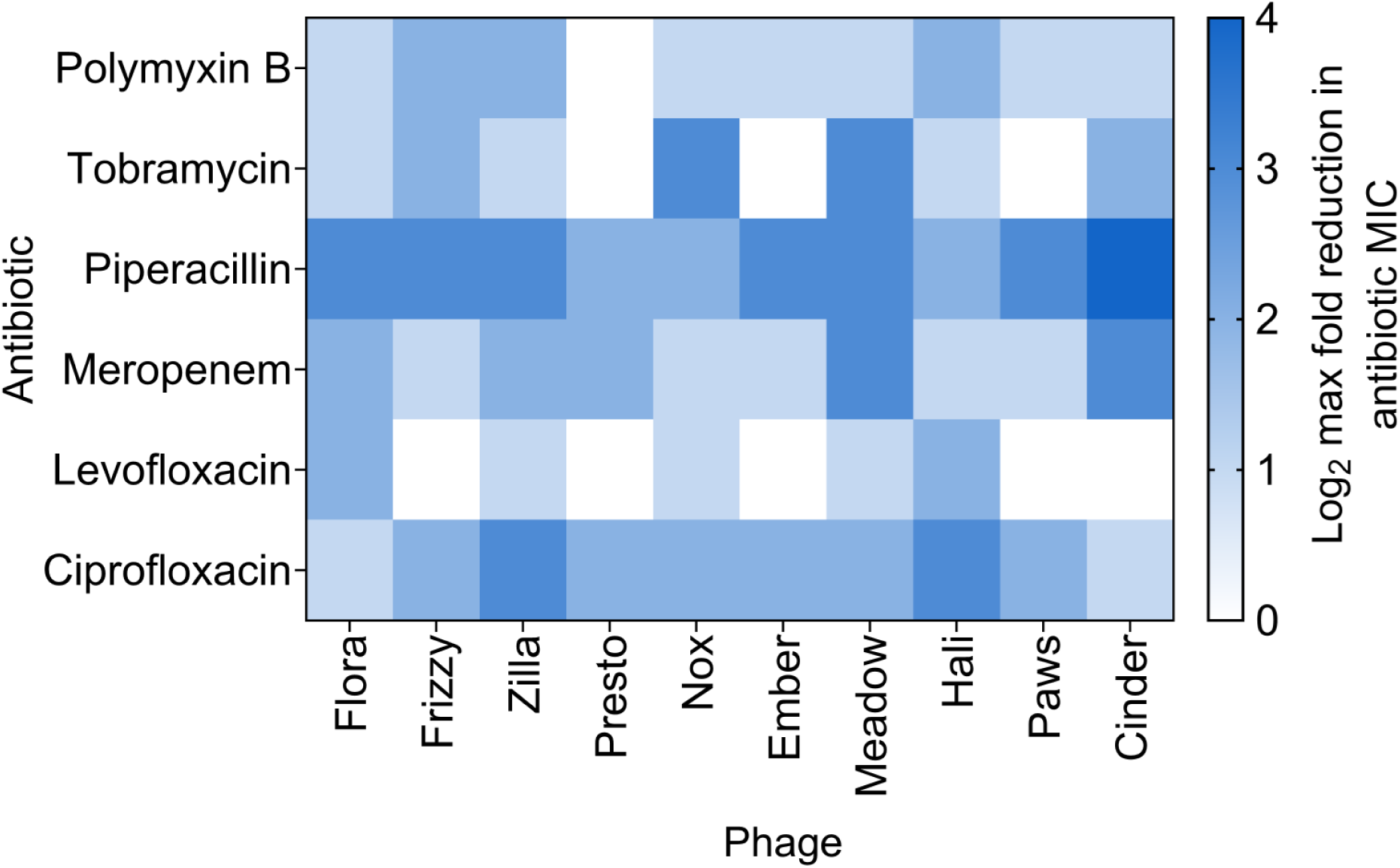
PAS is generalizable to other temperate phage-antibiotic pairings. Ten PA14 temperate phages were screened with two fluoroquinolones (ciprofloxacin and levofloxacin), two beta-lactams (meropenem and piperacillin), an aminoglycoside (tobramycin), and polymyxin B. Ciprofloxacin data are reproduced from Fig. 1c as a reference. Log_2_ maximum reduction in antibiotic MIC achieved in the presence of PA14 temperate phages relative to no phage control, plotted as a heat map (average of minimum n=3 biological replicates).

### Synergy with piperacillin reduces frequency of lysogeny, but not through induction

With a clear synergy observed with several antibiotics, we looked to confirm if the interaction with these other classes of antibiotics also operates at the level of biasing the phage lysis-lysogeny equilibrium. To test this, we purified survivors from our overnight survivor quantification assay performed for three other phage-antibiotic pairings. Phage Nox combined with tobramycin was able to synergistically reduce survivor counts by 2-3 logs, however there was no reduction in percent lysogeny observed, only reducing by 10% at the highest concentration tested (Fig. 6a). Meropenem combined with phage Meadow synergistically reduced survivor count but did not reduce frequency of lysogeny (Fig. 6b). In comparison, while there was not an impressive reduction observed in survivors for phage Cinder-piperacillin, there was a clear reduction in lysogeny from 60% for phage alone down to 25% in presence of sublethal piperacillin (Fig. 6c). The contrasting results of meropenem and piperacillin, belonging to the same drug class, further supports that tPAS is phage-antibiotic pairing specific.

**Figure 6.**
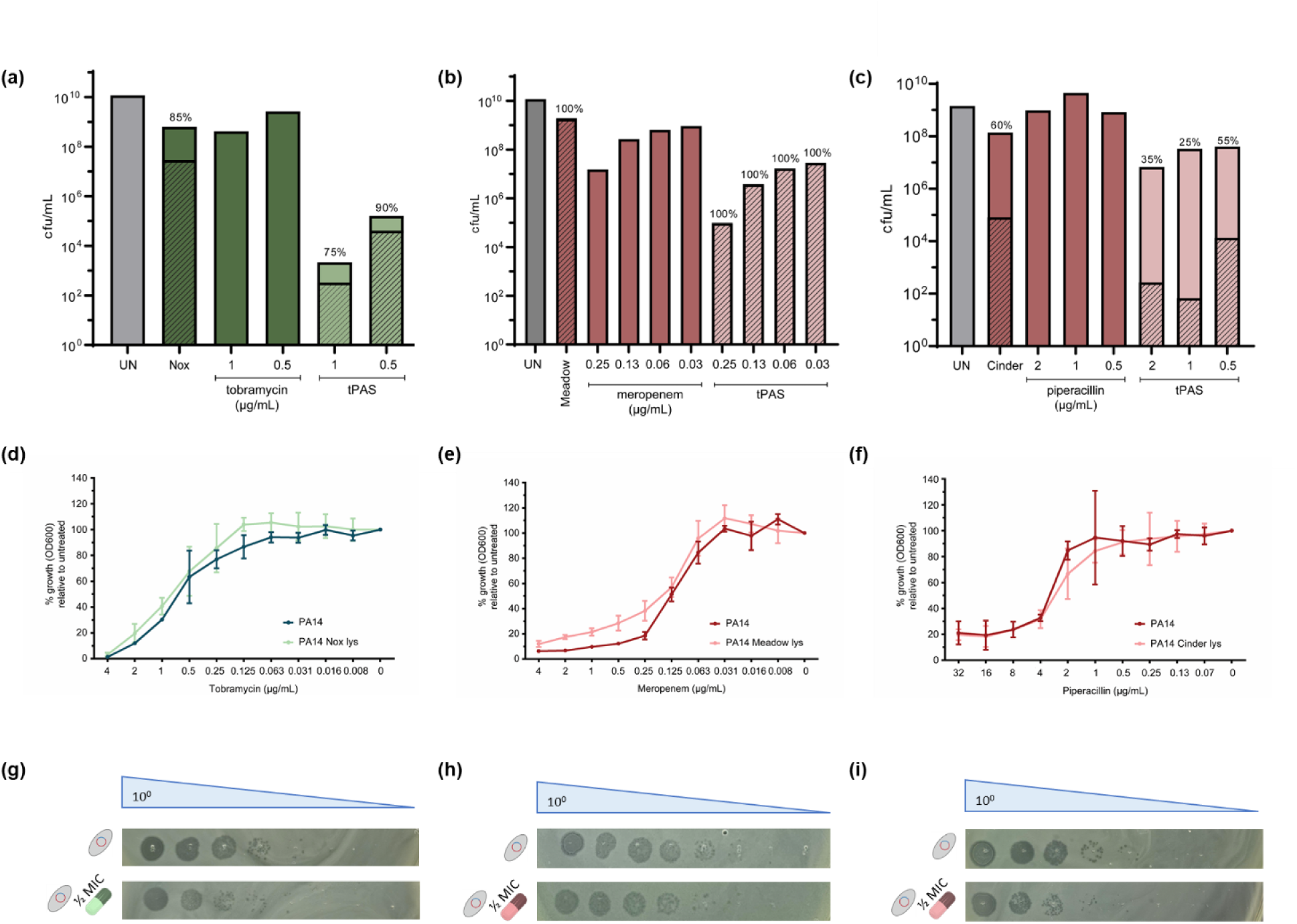
Piperacillin reduces frequency of lysogeny but not through induction. **(a-c)** Survivor quantification (cfu/mL) of PA14 challenged with individual and combination (a) phage Nox + tobramycin, (b) phage Meadow + meropenem, and (c) phage Cinder + piperacillin. Striped bars represent percentage of lysogens as determined using a lysogen stamp test, values also indicated above bar. Twenty survivors of phage ± antibiotic were streaked purified for lysogen testing. The height of the % lysogen bar is proportional to the height of the total cfu/mL bar (n=1 biological replicate). **(d-f)** Antibiotic sensitivity of wildtype PA14 and (d) phage Nox lysogen with tobramycin, (e) phage Meadow lysogen with meropenem, and (f) phage Cinder lysogen with piperacillin. MIC is represented as percent growth (endpoint OD600) relative to untreated host, plotted as mean ± SD (n=3 biological replicates, each in three technical triplicates). **(g-i)** Representative phage quantification ± ½ MIC antibiotic of PA14 (g) phage Nox lysogen with tobramycin, (h) phage Meadow lysogen with meropenem, and (i) phage Cinder lysogen with piperacillin (n=3 biological replicates, each with single technical replicate).

To confirm if the observed synergy and reduction in lysogeny could be a result of the ability of the antibiotic to result in induction, we tested the sensitivity of the wildtype and the lysogen for their respective antibiotic (Fig. 6d-i). None of the lysogens were more sensitive to the antibiotic or release more phages at sublethal doses relative to spontaneous induction after 18 h. This explains the lack of reduction in lysogeny observed with meropenem or tobramycin, where we hypothesize the synergy observed is likely driven by mechanism of traditional PAS instead. However, synergy with piperacillin does operate by biasing the lysis-lysogeny decision, however it is not at the level of induction. Piperacillin potentially works at the level of biasing the decision at the initial time of infection, like that reported with *E. coli* phage HK97 and gentamicin (43). The observed effect with piperacillin could also be attributed to its selective affinity for PBP3, unlike meropenem which preferentially binds PBP4 (64), and/or its ability to result in cell filamentation (65–67), where increase cell volume shows decrease probability of lysogeny in *E. coli* phage Lambda (68). This is the first report that we know of piperacillin’s ability to influence the phage lysis-lysogeny balance. Thus provides further proof that tPAS can be achieved with non-phage inducing antibiotics.

### tPAS is generalizable across clinical strains, even antibiotic-resistant ones

To establish whether the synergy is generalizable across hosts, we identified the six strongest phage-antibiotic pairings, one for each antibiotic in Fig. 6, and tested for synergy across multiple clinical strains, including both antibiotic sensitive and resistant ones. Fig. 6a shows maximum reduction in antibiotic MIC achieved with the addition of the phage across three isolates that were sensitive to all phages used. We also carried out this with five other clinical strains that were sensitive only to some overlapping phages (Supplementary Fig. 1).

Our findings show that the phage-antibiotic combinations that were initially identified to synergize in PA14 expand across multiple clinical isolates of *P. aeruginosa*, even in the multi-drug-resistant strain C0400. The reported ciprofloxacin resistance of this strain is consistent with our data, as the strain showed 4-fold higher MIC than PA14 (Supplementary Fig. 2a). We observed a reduction in ciprofloxacin MIC of this isolate even when combined with four other phages at levels much higher than those observed in the initial PA14 screen (Fig. 7b). The co-administration of temperate phage results in as high as an 8-fold reduction in MIC, bringing it down to levels comparable to PA14, re-sensitizing the isolate to the antibiotic.

**Figure 7.**
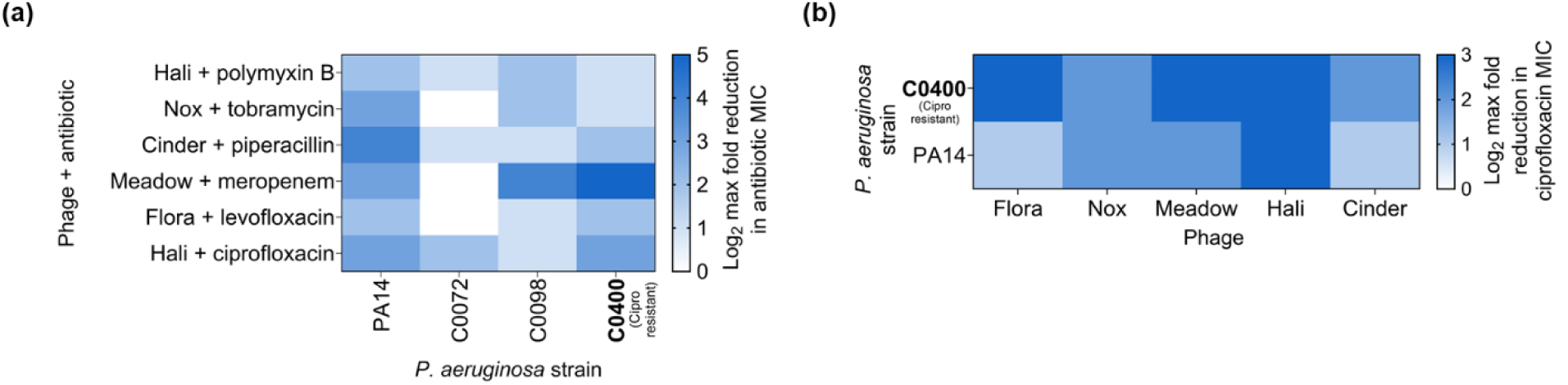
PAS with temperate phages works in clinical isolates. **(a)** Six of the strongest pairings from PA14 screen (phage Hali + ciprofloxacin, phage Flora + levofloxacin, phage Meadow + meropenem, phage Cinder + piperacillin, phage Nox + tobramycin, phage Hali + polymyxin B) were tested for synergy in *P. aeruginosa* clinical strains based on host range in Fig 2b. Result reported as log_2_ reduction in antibiotic MIC (n=3 biological replicates). **(b)** Log_2_ reduction in ciprofloxacin resistant strain C0400 MIC with the addition of five temperate phages. Average maximum reduction (n=3 biological replicates) in MIC shown as heat map as determined from checkerboard assays. Isolate in bold are classified as antibiotic-resistant strains in the clinical database using standard laboratory reporting.

Host C0400 is predicted by the Comprehensive Antibiotic Resistance Database Resistance Gene Identifier software to be ciprofloxacin resistant due to the presence of resistance-nodulation-cell division antibiotic efflux pump (69). Since many antibiotic targets also play a crucial role in phage replication and induction, the role of the exact mechanism of antibiotic resistance should be considered when investigating phage-antibiotic synergy. For example, efflux pumps can serve as phage receptors, hence development of phage resistance at the level of the surface receptor can result in re-sensitization to the antibiotic (70). This evolutionary tradeoff was used to successfully treat a 76-year-old male patient with *P. aeruginosa*-infected aortic graft using a combination of lytic phage OMKO1, which binds efflux pump, and ceftazidime.

However, the phages isolated in this study are temperate in nature where regrowth of lysogens present a major concern as opposed to surface receptor mutants. Despite originally being resistant to ciprofloxacin, host C0400 Hali lysogen displays a higher antibiotic sensitivity, compared to wild type C0400 and PA14 (Supplementary Fig. 2a) with significant increases in phage titer at a sublethal ciprofloxacin (Supplementary Fig. 2b). The increase in sensitivity of C0400 lysogen but lack of difference in PA14 lysogen ciprofloxacin sensitivity could potentially be explained by host specific factors. Since transposable phages are notorious for different integration sites, we also tested ciprofloxacin induction in independent lysogens, where our earlier findings suggest that it is the main phenotype driving tPAS with ciprofloxacin. We continued to detect ciprofloxacin triggered induction in other isolated C0400 Hali lysogens (not shown). These findings further support the hypothesis that Hali-ciprofloxacin synergy prevents lysogen overgrowth through antibiotic mediated induction, regardless of host. Our results highlight that tPAS can be achieved in clinical strains of *P. aeruginosa,* irrespective of their antibiotic sensitivity profile.

### Frequency of lysogeny and antibiotic mediated induction does not predict the ability of a temperate phage to synergize with an antibiotic

With proof that tPAS is working through biasing of the phage lysis-lysogeny equilibrium and a panel of phage-antibiotic pairings that showed efficacy in various hosts, we investigated if there are factors that can predict tPAS. We hypothesized that the frequency with which a phage lysogenizes its host or its susceptibility to induction with the antibiotic would serve as predictors of tPAS.

To determine if the ability of a phage to form stable lysogens in a specific host correlates with level of synergy, measured as reduction in MIC in the checkerboard assays, we infected host PA14 with seven phages and host C0400 with five phages in solid media and quantified frequency of lysogeny in the survivors of the phage challenge (Supplementary Fig. 4a). The frequency of lysogeny varied in PA14 ranging from as low as 26% for phage 12 to 100% for three other phages (phage Zilla, Drowsy, and Cinder). In comparison, all five phages exhibited 100% lysogeny in host C0400. However, there was no correlation between frequency of lysogenization and level of synergy observed with ciprofloxacin.

In addition, we also investigated the ability of sublethal ciprofloxacin to induce these prophages (Supplementary Fig. 4b). All tested lysogens of PA14 and C0400 were ciprofloxacin inducible, resulting on average in 10-1000-fold increase in phages released, except for the C0400 phage 1 lysogen. Similarly, when correlated with level of synergy as measured by decrease in MIC, level of ciprofloxacin mediated phage induction did not correlate with the magnitude of the synergy observed. Our results show that neither the frequency at which a phage forms lysogens or the extent to which the antibiotic results in induction correlates strongly with levels of synergy observed and therefore cannot predict if and how strongly a temperate phage will synergize with a particular antibiotic.

## Conclusion

Here we evaluate the generalizability of tPAS against *P. aeruginosa*. Temperate phages can be easily isolated from clinical strain collections and synergize with multiple clinically relevant antibiotics, irrespective of antibiotic target. This synergy can functionally lower antibiotic MICs up to 32-fold, and can do so even in resistant strains, functionally sensitizing them to the antibiotic. Mechanistically, some of the observed synergies are not temperate-phage specific, while others – notably ciprofloxacin, work at the level of induction in agreement with previous reports in a single phage in *E. coli* (42). Excitingly, piperacillin can also bias phage lysis-lysogeny equilibrium but appears to be doing so by biasing the initial lysis-lysogeny decision – in a manner akin to that reported for protein synthesis inhibitors for a single phage in *E. coli* (43). While temperate phages have been discarded in therapy due to their ability to lysogenize, the use of sublethal antibiotics can serve to bias away from lysogeny. This is both phage-host and phage-antibiotic pairing specific, and difficult to predict using factors inherent to the interaction between the two players.

## Supporting information

Phage Genome Sequences

## Acknowledgement

The authors would like to thank Gayatri Nair for helping to stamp the host range plates using the Singer HD Rotor. We would also like to thank Dr. Gerry Wright for access to the *P. aeruginosa* clinical isolates from the Wright Clinical Isolate Collection at the Institute for Infectious Disease Research (McMaster University, Ontario) and Dr. Lori Burrows for PA14.

R.F was supported by an NSERC Canadian Graduate Scholarship, Master’s and Doctoral scholarship. A.P.H. acknowledges funding through the Natural Sciences and Engineering Council of Canada (NSERC) Discovery Grant 2018-05996 and the Farncombe Family Chair in Phage Biology.

R.F. performed all the assays. A.P.H conceived the study. All authors contributed to the writing of the manuscript.

## Material and Methods

### Bacterial strains and growth conditions

*P. aeruginosa* strain PA14 was kindly gifted to us by the Burrows lab at McMaster University. Clinical strains of *P. aeruginosa* were obtained from the McMaster IIDR Wright clinical isolate collection. Bacterial strains were grown in 10 mL of lysogeny broth (LB) at 37°C with 130 rpm shaking (Ecotron, Infors HT, Quebec, Canada). For growth on solid media, 1% (w/v) of LB agar and 0.75% (w/v) of LB soft agar was used. All plates were incubated at 37°C from overnight growth.

For each experiment, same day culture was grown to an optical density (OD_600_) of 0.2 from a 1:100 dilution of an overnight culture. OD_600_ was measured using Thermo Fischer Scientific Spectronic 20D+ (Waltham, MA, USA).

### Phage isolation, propagation, and titration

One hundred and ninety-one *P. aeruginosa* clinical strains were grown overnight at 37°C in LB broth in a 96 well plate from frozen stocks. The plates were filtered using the Millipore MultiScreen_HTS_ vacuum manifold (Catalog MSVMHTS00, Darmstadt, Germany) with a Millipore Sigma MultiScreen_HTS_ High Volume 96-well 0.45 µm filter plate (Catalog MVHVN4525, Darmstadt, Germany). Approximately 2 µL of the undiluted filtrates were spotted on a 1% LB agar Nunc^TM^ Omnitray^TM^ single well plate with a 15 mL 0.75% agar overlay of containing 1 mL overnight culture of PA14. This assay would capture all antimicrobial components.

To identify phages, any filtrates that resulted in clearing were then confirmed by spotting serial dilutions where a phage would dilute to a single plaque as opposed to other bactericidal entities. Briefly, an agar overlay of 300 µL of PA14 overnight culture in to 3 mL of 0.75% molten agar was spread onto 1% LB agar petri plate. Ten-fold dilutions (10^0^ – 10^-7^) of the filtrates was prepared in 1x phage buffer and 3 µL was spotted on overlay.

The phages were amplified in LB broth to increase the titer. Primary amplification was carried out using frozen stock of the bacteria and phage inoculated in 10 mL of LB broth and incubated for 18 h. Secondary amplification was carried out by challenging same day grown culture at OD_600_ 0.2 with 50 µL of the primary amplification. Phages were titered on the respective host using a spot test and standard plaque assay.

To determine multiplicity of infection (MOI), an OD_600_ vs colony forming unit (cfu/mL) standard curve was carried out for each bacterial strain. Optical density of a same day culture was measured at 30 min intervals and 50 µL of culture was sampled at every hour, diluted 10-fold in LB, and 100 µL was plated on 1% LB plate using glass beads. MOI was calculated as 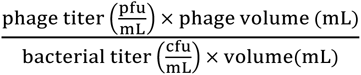.

For the clinical isolate screen, phages were amplified on the host being tested if the efficiency of plaquing was 2 log_10_ or lower relative to PA14. If not, lysate prepared on PA14 was used for the checkerboard screens.

### Minimum inhibitory concentration

The minimum inhibitory concentration of each antibiotic was determined using a modified microtiter assay. Ciprofloxacin (hydrochloride) was obtained from Cayman Chemicals (Catalog 14286-5, Ann Arbor, Michigan, USA), levofloxacin from Cedarlane (Catalog 20382-1, Burlington, ON, Canada), meropenem from Sigma-Aldrich (Catalog PHR1772, Oakville, ON, Canada), piperacillin from Sigma-Aldrich (Catalog PHR1805, Oakville, ON, Canada), tobramycin from Sigma-Aldrich (Catalog PHR1079, Oakville, ON, Canada), and polymyxin B sulfate from EMD Millipore (Catalog D46530, Oakville, ON, Canada). In a narrow 96 well plate (Corning, Product Number 3370, ME, USA), 100 µL of same day culture was combined with antibiotic stock and nuclease water in a final volume of 250 µL. The plate was incubated for 18 h at 37°C with no agitation. The endpoint OD_600_ was measured using the Epoch 2 microplate spectrophotometer (BioTek Instruments, Inc., VT, USA) with a 10 sec double orbital shake before read. MIC was the lowest antibiotic concentration that resulted in no growth and was re-evaluated for every new batch of antibiotic stock prepared.

### Checkerboard assay

Same day culture grown to OD 0.2 was challenged with two-fold dilutions of the phage on the vertical axis and two-fold dilution of the antibiotic on the horizontal axis in a narrow 96-well plate. MOI tested for each phage was 40 – 1.25. The antibiotic concentrations tested changed depending on the MIC. For each checkerboard performed, the culture, phage, and antibiotic volume were fixed to 25 µL, 62.5 µL, and 12.5 µL, respectively, to achieve the desired MOI and antibiotic concentration. Untreated host and LB-only were used as growth controls. Plates were incubated for 18-20 h at 37°C. OD_600_ was used to measure growth in each plate using a BioTek Epoch 2 microplate spectrophotometer. The percent growth in each well relative to untreated host control was used to determine the maximum reduction in antibiotic MIC that was achieved with the addition of the phage, regardless of MOI.

### Phage genome sequencing and analysis

Phage DNA extraction, DNA library preparation, sequencing and analyzes were carried out as described in (71). Briefly, 500 µL of phage lysate (≥ 10^8^ pfu/mL) was treated with DNase I, RNase, and DNase I reaction buffer and incubated at 37°C for 30 min followed by DNase and RNase inactivation at 65°C for 10 min. Next, proteinase K and 2% final volume SDS was added and incubated for 1 h at 37°C to denature protein and break open phage capsid. Following incubation, mixture was split into two aliquots and equal volume of phenol-chloroform was added. The mixture was centrifuged for 10 min at >13 rpm and supernatant was collected and treated with 1/10^th^ vol of ammonium acetate and one volume of −20°C isopropanol, followed by another round of 10 min centrifugation. The pellet was resuspended in −20°C 70% ethanol and centrifuged for 10 min. After decanting the supernatant, the new pellet was air dried for 15 min and rehydrated in elution buffer overnight at 4°C. The DNA concentration was measured using Invitrogen Qubit 4 Fluorometer.

DNA libraries were prepared for sequencing using the NEBNext® Ultra™ II DNA Library Prep Kit (New England Biolabs, catalog no. E7645S, Massachusetts, USA) with a modified version of the Derakhshani *et. al.*, (2020) protocol as outlined in (71). Sequencing was carried out using MiSeq with paired-end 2 x 300 reads at the McMaster Metagenomics Facility (Ontario, Canada). Genome assembly and analysis was carried out using the pipeline outlined in (71). Briefly, quality assessment on raw reads was carried out using FastQC v0.11.8 (Andrews, 2010) before and after trimming with Trimmomatic v0.38 (Bolger *et al.,* 2014). The trimmed reads next go through a series of steps in which *de novo* assembly is performed using metaSPades v3.13.0 (Meleshko, 2017) and the sequence is predicted as phage or not. The last step in the pipeline involves analysis which includes within sample comparison to determine how similar samples are to each other and genome annotation using RASTtk v1.3.0 (Brettin, 2019). The tree was generated using VipTree version 4.0 (https://www.genome.jp/viptree/) using the default setting after which a smaller subset of phages was selected to regenerate a smaller tree.

### Host range analysis

Phage host range was carried out as described in (71). Briefly, a micro plaque assay was carried out using the Singer Rotor HDA. Ninety-eight clinical isolates of *P. aeruginosa* were inoculated from frozen in 1 mL LB in a deep 96 well plate and grown overnight. Forty-five microliters of each isolate were transferred into a 384-well plate such that each of the four quadrants of the 384 well plate are replicates. One 384-well plate was prepared for each phage where each quadrant contains 45 µL of either undiluted, 10^-2^, 10^-4^, and 10^-6^ phage lysate dilution prepared in LB broth. Stamping plates were prepared with 25 mL of 1.5% LB agar per plate. Using the Singer Rotor HDA, culture was stamped first, followed by phage directly on top at a 1536 density (four replicates per phage dilution-host pairing). One plate with only culture stamped was used as a control. The plates were wrapped in plastic bags and incubated for 18 h at 37°C. Phage sensitivity and efficiency of plaquing, the lowest dilution at which plaques were observed, was noted for each phage-host pairing.

### Survivor quantification assay

Overnight challenge survivor quantification assay was performed as previously described in (42). Briefly, same day culture of PA14 was challenged with phage alone at an MOI of at least 10, two-fold dilutions of antibiotic alone starting at MIC, and in combination in a final volume of 1 mL. Untreated host was used as a growth control. Challenges were incubated at 37°C with 130 rpm shaking for 18 h, following which 100 µL of 10-fold dilutions carried out with LB broth was added to 5 ml of LB soft agar and spread onto an empty petri plate. Survivors were counted after an overnight incubation to determine the fold reduction in survivor count relative to the untreated host. The expected synergistic effect was calculated by multiplying phage alone reduction with the appropriate antibiotic alone challenge. For frequency of lysogenization characterization, twenty survivors for phage and all phage + antibiotic challenged were streaked purified thrice on 1% LB and lysogen detection was performed as described below.

### Overnight challenge phage quantification

For each survivor quantification assay performed, half the volume after the 18 h challenge was filtered using a Millipore MultiScreen_HTS_ vacuum manifold (Catalog MSVMHTS00, Darmstadt, Germany) vacuum manifold and Millipore Sigma MultiScreen_HTS_ High Volume 96-well 0.45-micron filter plate (Catalog MVHVN4525, Darmstadt, Germany). Phages in the filtrates were quantified using a standard plaque assay using PA14 as the host.

### Lysogen isolation & detection

Spot tests of phage 1, 3, 5, 8, 12, 13, and 17 were carried out on PA14 and phage 1, 5, 12, 13, and 17 on *P. aeruginosa* clinical strain C0400. All lysates were previously amplified on host PA14 in broth. Regrowth on the plate from the phage challenge was streaked out on to 1% LB agar plate. Twenty colonies for each phage-host pair were streak purified three times and inoculated in 1mL LB broth overnight in a deep 96 well plate. Wild type culture, not exposed to the phages, and LB broth were added as control. The plates were incubated for 18-24 h at 37°C with no agitation. The following day, the cultures were stamped on an agar overlay of either PA14 or C0400, depending on the original host used for the spot test, using a disposable pin replicator. Cultures that resulted in the clearing of the wildtype host were classified as lysogen. Frequency of lysogeny was calculated as percent of the total number of survivors that were characterized as lysogens for each phage-host pairing. One lysogen for each phage-host pair was randomly selected for ciprofloxacin induction.

### Ciprofloxacin induction

Same-day culture of wildtype host and the lysogen were grown in the presence of two-fold dilutions of ciprofloxacin, prepared in nuclease free water, in a narrow 96 well plate in 250 µL final volume. Nuclease free water was added to the no antibiotic growth control. The plates were incubated for 18 h at 37°C with no agitation. The following day, the plate was filtered using a vacuum manifold with a 0.45-micron filter plate. The phages in the filtrate of the untreated host (wild type and lysogen), and the host treated with antibiotic at MIC, ½ MIC, and ¼ MIC of that specific trial were quantified using a spot test.

### Delay administration

Same-day PA14 culture was treated with phage Hali amplified on PA14 (MOI at least 10), ciprofloxacin (2-fold dilution MIC – 1/16 MIC), and combination phage Hali + ciprofloxacin in a final volume of 250 µL in a narrow 96 well plate. To test the effects of delayed administration, either antibiotic or phage was withheld for 4 h. Co-administration was tested on the same plate for each delayed administration tested. Each condition was set up in triplicates. LB broth was used as a negative control. Growth curves were carried out using OD_600_ for 4 h at 30 min intervals using the Epoch 2 microplate spectrophotometer. For delayed antibiotic, no treatment control and phage challenge were treated with nuclease free water. For delayed phage, LB broth was added to the no treatment and antibiotic alone challenge. After the addition of the delayed agent, growth in the plate was measured for another 18 h at 30 min intervals. Endpoint growth (OD600) after 18 h was calculated relative to untreated culture and plotted as a heatmap.

## Supplementary

**SUPPLEMENTARY Figure 1.**
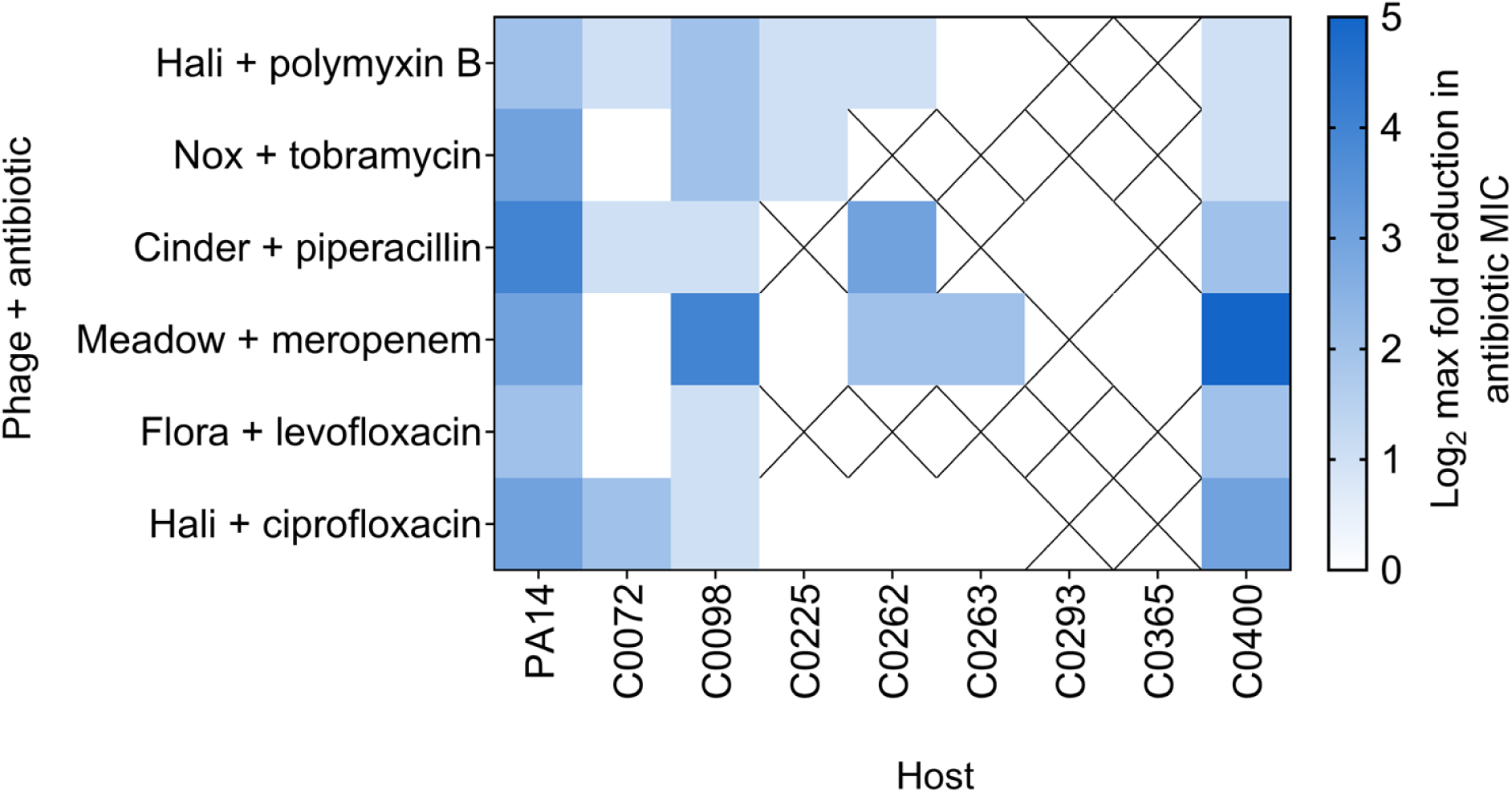
PAS with temperate phages across clinical isolates. Reduction in antibiotic MIC of multiple clinical strains with the addition of temperate phage. Average maximum reduction (n=3 biological replicates) in MIC shown as heat map as determined from checkerboard assays. “X” denotes pairings that were not tested in the specified strain either due to lack of phage sensitivity or inability to obtain a high enough titer for checkerboard assay.

**SUPPLEMENTARY Figure 2.**
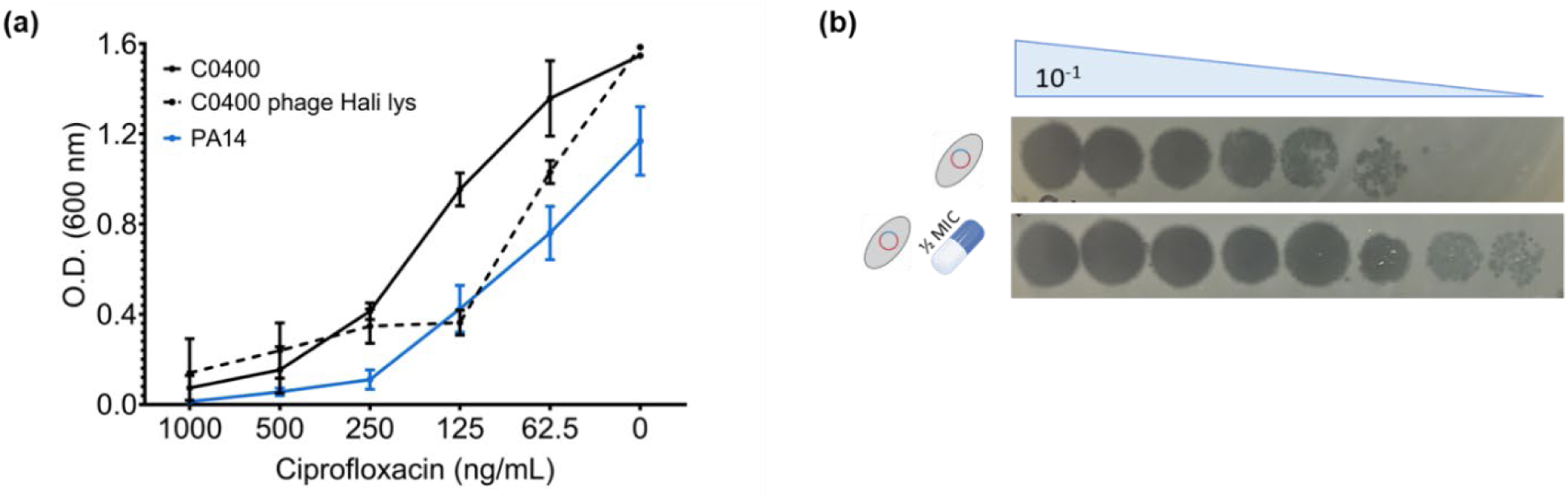
C0400 phage Hali lysogen is ciprofloxacin inducible. **(a)** End point growth (OD600) of C0400 (black line), C0400 phage Hali lysogen (dotted black line), and PA14 (blue line) across a range of ciprofloxacin concentrations (µg/mL) after 18 h. Data shown as mean ± SD (n=3 biological replicates, each in technical replicates). **(b)** Representative end point phage titer of filtrates of host C0400 and C0400 phage Hali lysogen no or ½ MIC ciprofloxacin treatment.

**SUPPLEMENTARY Figure 3.**
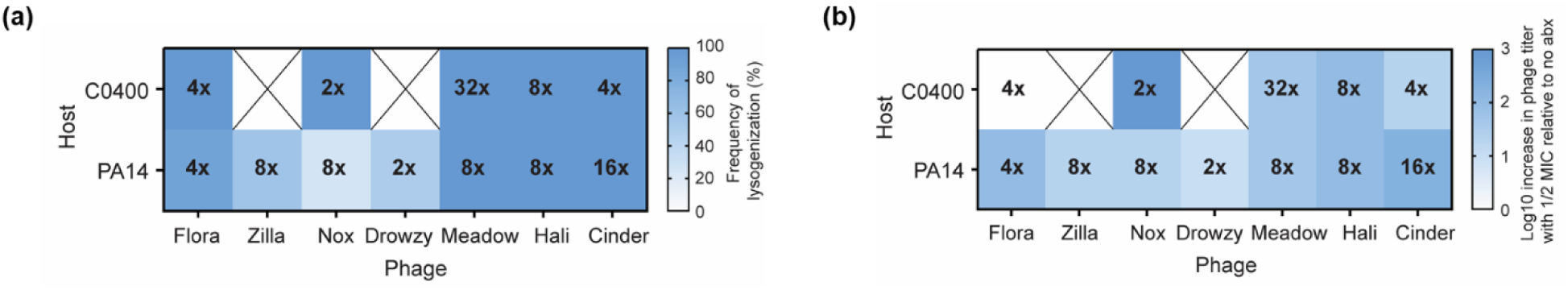
Level of synergy does not correlate with frequency of lysogenization or level of induction. **(a)** Frequency of lysogenization temperate phages in host PA14 and C0400 as a heat map. Twenty survivors from phage challenge on solid media were purified and screened for the presence of the phage using a streak test. Number within the colored box indicate the reduction in ciprofloxacin MIC observed in checkerboard assay. **(b)** Log10 average increase (n=3 biological replicate, each in single technical replicate) in phage titer when PA14 or C0400 lysogen challenged with ½ MIC ciprofloxacin relative to no antibiotic control, represented as a heat map. X denotes combination not tested.

